# 3D Shape Modeling for Cell Nuclear Morphological Analysis and Classification

**DOI:** 10.1101/313411

**Authors:** Alexandr A. Kalinin, Ari Allyn-Feuer, Alex Ade, Gordon-Victor Fon, Walter Meixner, David Dilworth, Syed S. Husain, Jeffrey R. de Wet, Gerald A. Higgins, Gen Zheng, Amy Creekmore, John W. Wiley, James E. Verdone, Robert W. Veltri, Kenneth J. Pienta, Donald S. Coffey, Brian D. Athey, Ivo D. Dinov

**Affiliations:** Department of Computational Medicine and Bioinformatics, University of Michigan Medical School, Ann Arbor, MI, USA; Statistics Online Computational Resource (SOCR), Department of Health Behavior and Biological Sciences, University of Michigan School of Nursing, Ann Arbor, MI, USA; Division of Gastroenterology, Department of Internal Medicine, University of Michigan Medical School, Ann Arbor, MI, USA; Department of Urology, James Buchanan Brady Urological Institute, Johns Hopkins University School of Medicine, Baltimore, MD, USA; Michigan Institute for Data Science (MIDAS), University of Michigan, Ann Arbor, MI, USA

## Abstract

Quantitative analysis of morphological changes in a cell nucleus is important for the understanding of nuclear architecture and its relationship with pathological conditions such as cancer. However, dimensionality of imaging data, together with a great variability of nuclear shapes, presents challenges for 3D morphological analysis. Thus, there is a compelling need for robust 3D nuclear morphometric techniques to carry out population-wide analysis. We propose a new approach that combines modeling, analysis, and interpretation of morphometric characteristics of cell nuclei and nucleoli in 3D. We used robust surface reconstruction that allows accurate approximation of 3D object boundary. Then, we computed geometric morphological measures characterizing the form of cell nuclei and nucleoli. Using these features, we compared over 450 nuclei with about 1,000 nucleoli of epithelial and mesenchymal prostate cancer cells, as well as 1,000 nuclei with over 2,000 nucleoli from serum-starved and proliferating fibroblast cells. Classification of sets of 9 and 15 cells achieved accuracy of 95.4% and 98%, respectively, for prostate cancer cells, and 95% and 98% for fibroblast cells. To our knowledge, this is the first attempt to combine these methods for 3D nuclear shape modeling and morphometry into a highly parallel pipeline workflow for morphometric analysis of thousands of nuclei and nucleoli in 3D.

## Introduction

### Motivation

Cell nuclear morphology is regulated by complex underlying biological mechanisms related to cell differentiation, development, proliferation, and disease^1–3^. Changes in the nuclear form are associated with reorganization of chromatin architecture related to altered functional properties such as gene regulation and expression^1,3^. Moreover, many studies in mechanobiology show that geometric constraints and mechanical forces applied to a cell deform it and, conversely, affect nuclear and chromatin dynamics, as well as gene and pathway activation^4,5^. Thus, nuclear morphological quantification becomes of major relevance as studies of the reorganization of the chromatin and DNA architecture in the spatial and temporal framework, known as the 4D nucleome, emerge^6,7^. Cellular structures of interest in the context of the 4D nucleome include not only the nucleus itself, but also the nucleolus and nucleolar-associating domains, chromosome territories, topologically associating domains, lamina-associating domains, and loop domains in transcription factories^6,8^. Furthermore, understanding these processes through quantitative analysis of morphological changes also has many medical implications, for example, in detection, understanding, and treatment of pathological conditions such as cancer^7–10^.

While efforts have been made to develop cell and nuclear shape characteristics in 2D or pseudo3D^11,12^, several studies have demonstrated that 3D morphometric measures provide better results for nuclear shape description and discrimination^13–15^. However, 3D shape descriptors that permit robust morphological analysis and facilitate human interpretation are still under active investigation^16^. Additionally, the dimensionality and volume of acquired data, various image acquisition conditions, and great variability of cell shapes in a population present challenges for 3D shape analysis methods that should be scalable, robust to noise, and specific enough across cell populations at the same time. Thus, there is a compelling need for robust 3D nuclear morphometric techniques to carry out population-wide analysis^17^.

### 3D shape representation and morphometric measures

The way cell nuclear shapes can be measured depends on their representation extracted from image data^11^. Many 3D morphometric measures are applied “as is” to 3D geometric objects represented by volumetric data^18^. However, voxels-based shape representations are noisy, and they may lose fine geometric details or even break the object’s topological structure. Moreover, these representations are not intrinsic, and vary when changing pose or deforming the object. A recent review of approaches to 3D cell shape description^16^ separated them into three categories in increasing order of complexity: landmark-based, graph-based, and moment-based. This last category includes approaches that are widely used in cellular morphology and allow the user to obtain a global representation that combines low-order moments describing the coarse conformation with high-order moments retaining information at higher frequency. Typically, before applying these methods, a binary mask or outline of the shape (surface) is first extracted from image data, which is done by most segmentation methods. These masks are assumed to have a sphere-like topology and can be projected onto an appropriate basis. Two popular approaches of this type are spherical harmonics (SPHARM)^19^ and spherical wavelets^20^. Both methods first map the surface of interest onto the sphere using appropriate spherical parameterization techniques, and then project it onto a reference function basis living on the sphere. SPHARM is arguably one of the most widely applied cell morphology modeling approaches^21–24^. In SPHARM the spherical signal is projected onto a basis of Legendre polynomials, extending the classical Fourier analysis to signals on the two-sphere. SPHARM coefficients describe general conformation of the shape of interest at different spatial scales, are rotation invariant, and can be directly used as features for further analysis^25^. However, SPHARM methods are most appropriate when low order approximation is satisfactory and become less effective in preserving surface details, as artificial oscillations start to appear when higher order basis functions are incorporated^26^. More robust smooth surface reconstruction can be obtained from a 3D binary mask via Laplace-Beltrami (LB) eigen-projection, followed by topology-preserving boundary deformation to remove various artifacts^26^. On a unit sphere, the LB eigen-functions correspond to spherical harmonics, so overall they can be viewed as a generalization of the SPHARM to the complex geometry manifold with local adaptation of the basis to the dataset at hand^27^. The proposed method has been demonstrated to produce smoother and more detailed surfaces compared to the SPHARM and the topology preserving level sets^28^. Extracted surfaces are smooth, accurately represent the shape of an object, and can be further used for morphometric analysis.

In order to extract shape geometric characteristics, boundary surfaces of binary masks are typically reconstructed from voxel data and discretized as meshes. At the next step, various useful morphometric descriptors can be computed based on this representation. Useful extrinsic and intrinsic geometric descriptors aim to distinguish between global and local shape features. Intrinsic measures capture shape properties that are invariant under transformations (e.g., affine: rotation, translation and scaling). Various shape morphometry measures, like surface area and Gaussian curvature, represent invariant metrics of complexity, which are stable under special transformations of the surface (e.g., bending) that do not affect the inner geometry of the boundary of the 3D volume^29^. Alternatively, shape metrics, e.g., mean *L*^2^-norm and the extrinsic curvature index, are sensitive to affine transformation and other shape morphology in the ambient space. Shape index and curvedness are morphometric descriptors that can capture local shape features, independently or in relation to the size of an object^30^. Combination of the object surface reconstruction with the extraction of such shape measures demonstrated high performance in recent neuroimaging studies for discriminatory morphometric analysis of complex 3D shapes of cortical and subcortical brain areas^31–33^.

### Technical capabilities and interoperability of tools

When it comes to a choice of tools for 3D cell nuclear morphometrics, reproducibility and implementation availability are among major concerns in the field of bioimage analysis^16^. To date, many of the widely available software tools for cell shape morphometry were either developed for the analysis of 2D^11,34–38^ or pseudo-3D images^39^. Other tools only implement slice-by-slice or voxel-based morphometry^40–43^, providing a coarse approximation of the global cell shape that is sensitive to increasing amounts of noise and usually fails to characterize morphological variations occurring at different spatial scales. Other common limitations of many 3D cell morphology solutions include a lack of high-throughput processing capabilities or restrictions to the specific programming language or platform that dictate principles of tool implementation^44–46^. Implementations of methods in a bioimage analysis landscape are highly diverse. They range across programming languages, software libraries, and file formats, which increases module interoperability issues and makes code reuse extremely difficult. Re-implementing underlying methods is often very challenging, time-consuming, and error prone^47^. Some of the existing bioimage analysis frameworks, including ImageJ^48^, rely on a plugin architecture, which allows their extension via third-party contributions^40,41,43^. High-throughput capabilities of some of these software packages are limited to processing of multiple objects simultaneously within its graphical user interface (GUI), for example, Tango^43^. More advanced packages, such as CellProfiler 2.0^37^, BioimageXD^42^, and Icy^41^, provide a basic graphical interface to assemble elementary tasks into reusable pipelines that make it possible to execute in GUI and batch modes. However, these solutions are still limited to specific scripting languages and libraries supported by the main software package. They also don’t provide a straightforward way to take advantage of the growing number of parallel hardware configurations, such as clusters, clouds, and high-performance computing, which limits the scalability of these solutions.

An alternative to plugin-based solutions, software platforms with modular design allow integration of already existing solutions into workflows without re-implementing them in a specific language, and provide methods for optimizing module interaction, re-usage, and extension^49^. An example of an extensive and feature rich solution for building and executing complex workflows is the LONI Pipeline^31,50^. This client-server platform enables users to efficiently describe atomic modules and end-to-end protocols in a graphical canvas using a large library of powerful computational tools. The Pipeline back-end server has extensive support for parallel execution on a grid cluster, including automated data converting, formatting and transfer, optimal job submission and management, pausing execution, and combining local and remote software and data sources. Most importantly, it makes it very easy to create new custom modules from any software that supports a command line interface (CLI). The Pipeline allows users to take advantage of a highly diverse set of tools and connect them together as steps of a computational protocol that is then executed in a high-throughput, parallel fashion. Validated individual modules and end-to-end workflows may be saved, reused in other workflows, easily modified and repurposed. Additionally, the LONI Pipeline saves information about executed steps (such as software origin, version, and architecture) providing provenance information^50,51^.

### Study aims

This study has two complementary aims. The *first aim* is to assess and validate 3D morphometry metrics for nuclear and nucleolar shape description and classification. Improving the discriminative performance in terms of statistical metrics has been the driving force behind our methodological efforts and selection of specific tools in this work. First, surfaces of 3D masks extracted from the microscopy data are reconstructed using Laplace-Beltrami eigen-projection and topology-preserving boundary deformation^26^. Then, we computed intrinsic and extrinsic geometric metrics, which are used as derived signature vectors (shape biomarkers) to characterize the complexity of the 3D shapes and discriminate between observed clinical and phenotypic traits. These metrics include volume, surface area, mean curvature, curvedness, shape index, and fractal dimension^30,52,53^. Although these methods were previously used in recent neuroimaging studies^31–33^, this is the first attempt, to our knowledge, to apply robust smooth LB-based surface reconstruction with intrinsic and extrinsic morphometric measure extraction to 3D cell nuclear and nucleolar shape modeling and morphometry. Suggested modeling and analysis methods are not restricted to nuclear and nucleolar shapes and can be used for the shape quantification of other cellular compartments, depending on their topology.

The *second aim* is to develop a reproducible pipeline workflow implementing the entire process that can be customized and expanded for deep exploration of associations between 3D nuclear and nucleolar shape phenotypes in health and disease. High-throughput imaging (HTI) can include automatization of liquid handling, microscopy-based image acquisition, image processing, and statistical data analysis^17^. Our work focuses on the last two aspects of this definition. We implemented a streamlined multi-step protocol using a diverse set of tools to achieve optimal performance compared to alternatives at each step of analysis. These tools are represented as individual modules seamlessly connected in the LONI Pipeline workflow. This workflow meets modern standards for high-throughput imaging processing and analysis and is mostly automated with a focus on validity and reproducibility. Our implementation is massively parallel, customizable, and provides fully automated execution and data provenance out-of-the-box. At the final step of the workflow, we employed machine learning methods to investigate the associations between cell phenotypes and treatment conditions using cell shape morphometric measures as features. We show that using a combination of 3D nuclear and nucleolar morphometry improves the discrimination between *in vitro* cell conditions of human fibroblast and human prostate cancer (PC3) cell lines.

To promote the reproducibility of results, facilitate open-scientific development, and enable collaborative validation, we made the pipeline workflows, together with underlying source code, documentation, and derived data from this study, available online^54^. The workflow will be made available via the LONI Pipeline, along with publicly available computational resources to showcase an online demonstration.

## Methods

Fig. 1 shows a high-level view of the end-to-end protocol. We start with a dataset of 3D binary nuclear and nucleolar masks. We modeled 3D nuclear and nucleolar boundaries by their surface reconstruction and extracted the derived morphometry measures. Finally, we computed statistical differences, identified shape morphometry-phenotype associations, and evaluated the results.

**Figure 1.**
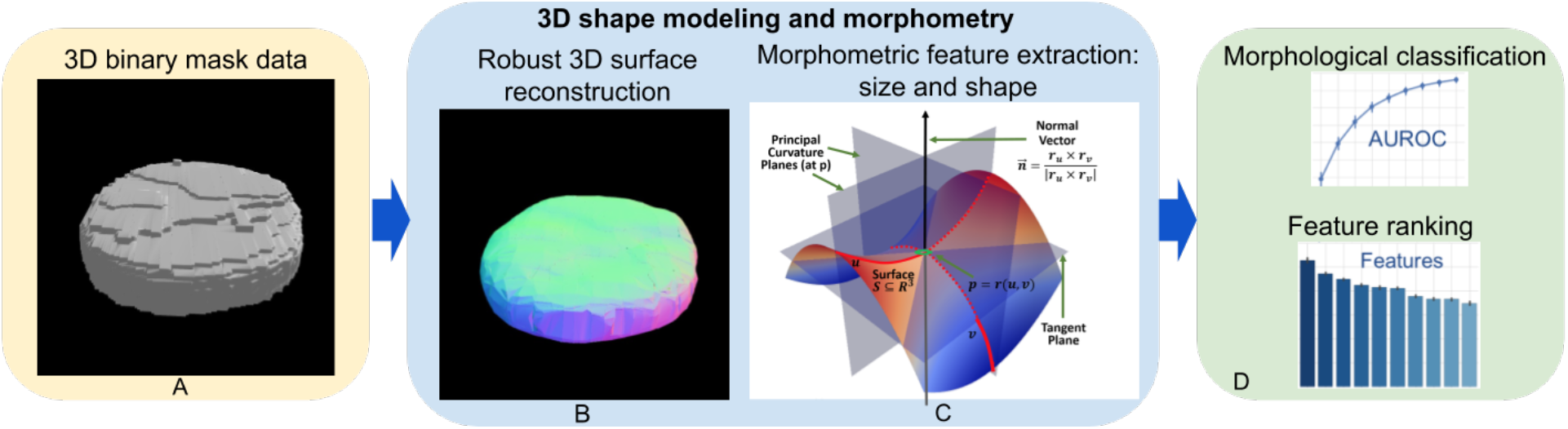
High-level schematic flow of the 3D image processing protocol: (A) 3D binary mask data; (B) mathematical representation and modeling of shape and size; (C) calculation of derived intrinsic and extrinsic geometric measures; and (D) machine learning based classification, feature ranking, and analysis.

### Dataset description

In this study we used the 3D Cell Nuclear Morphology Microscopy Imaging Dataset, one of the largest publicly available 3D cell imaging datasets to date^18^. This dataset consists of two collections of 3D volumetric microscopic cell images with corresponding nuclear and nucleolar binary masks. Each collection includes images of cells in two phenotypic states, and thus poses a binary classification problem with image-level labels that can be used for the assessment of cell nuclear and nucleolar morphometric analysis. Binary masks in each collection are obtained by segmentation of the original data. Nuclear masks are extracted from a DAPI (4',6-diamidino-2-phenylindole) channel, while fibrillarin antibody-stained (anti-fibrillarin) and ethidium bromide-stained (EtBr) channels are both used for nucleolar binary mask extraction (see^18^ for details). Segmented binary masks are represented by 1024×1024×Z 3D TIFF sub-volumes. For every mask sub-volume, accompanying vendor metadata extracted from the original data are available for analysis, as well.

### Robust smooth surface reconstruction

To model the 3D shape of cell nuclei and nucleoli, boundaries of their 3D masks extracted from the microscopy data are modeled as genus zero two-dimensional manifolds (homeomorphic to a 2-sphere *S*^2^)^55^ that are embedded as triangulated surfaces in ℝ^3^, Fig. 1B. Our approach uses an iterative Laplace-Beltrami eigen-projection and a topology-preserving boundary deformation algorithm^26^. This algorithm performs robust reconstruction of the objects’ surfaces from their segmented masks using an iterative mask filtering process. First, a mesh representation is constructed from the boundary of an object’s binary mask. Then, the boundary is projected onto the subspace of its Laplace–Beltrami eigen-functions^27^, which allows the algorithm to automatically locate the position of spurious features by computing the metric distortion in eigen-projection. LB eigen-functions are intrinsically defined and can be easily computed from the boundary surface with no need of any parameterizations. They are also isometry invariant, and thus robust to the jagged nature of the boundary surface, which is desirable for biomedical shape analysis^56^. In our prior experience^26^, the discretized LB spectrum captures intrinsic shape characteristics (e.g., global shape transformations will preserve the spectral signature). The magnitude of the eigenvalues of the LB operator intuitively corresponds to the frequency in Fourier analysis, thus it provides a convenient mechanism to control the smoothness of the reconstructed surface. Using this information, the second step is a mask deformation process that only removes the spurious features while keeping the rest of the mask intact, thus preventing unintended volume shrinkage. This deformation is topology-preserving and well-composed such that the boundary surface of the mask is a manifold. The last two steps iterate until convergence and the method generates the final surface as the eigen-projection of the mask boundary, which is a smooth surface with genus zero topology^26^. These properties allow application of this algorithm to any shape, including, for example, crescent-shaped, multi-lobed, and folded, as long as shape topology is homeomorphic to a sphere. The exemplar results of this step performed on nuclear and nucleolar masks are shown in Fig. 2.

**Figure 2.**
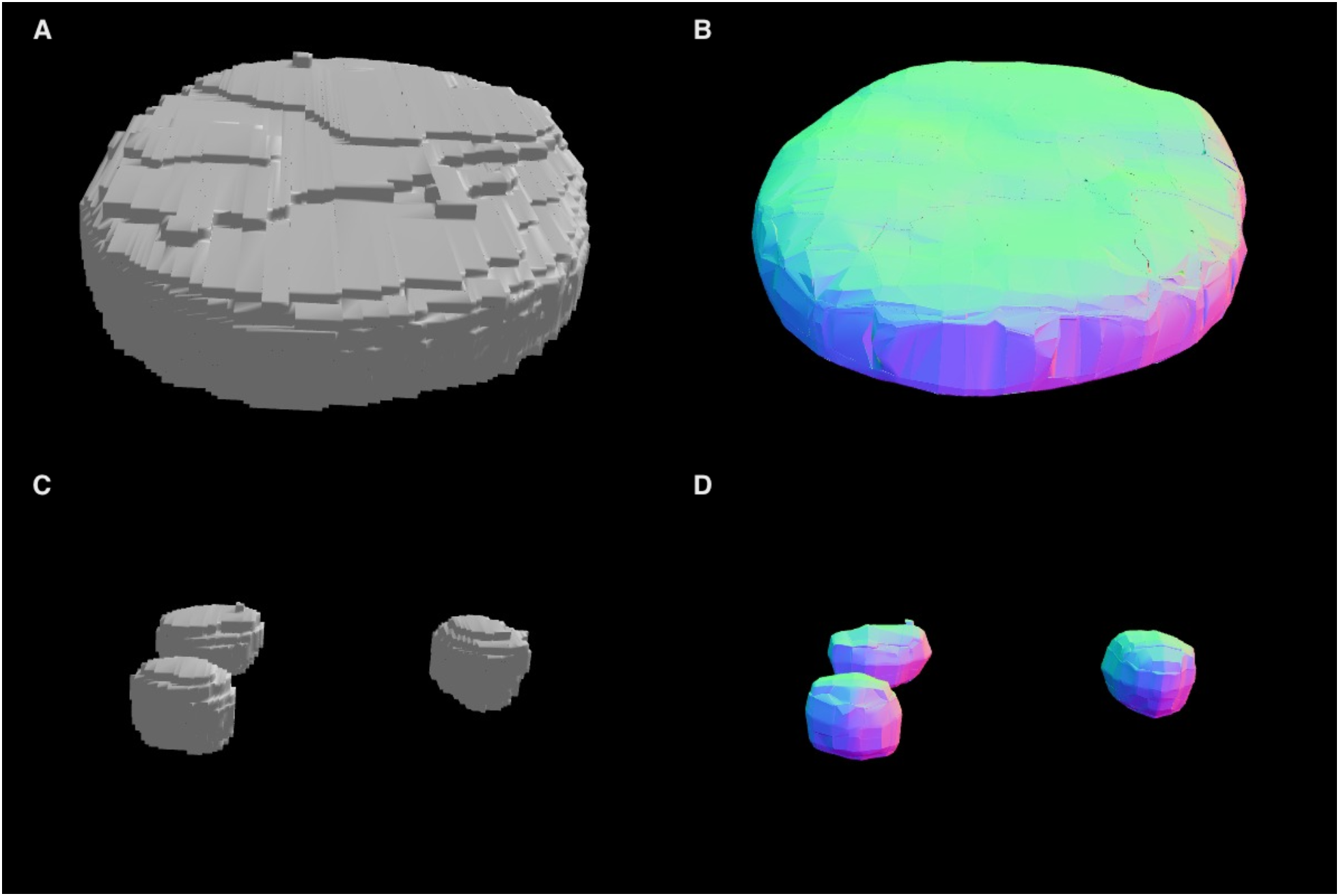
Robust smooth surface reconstruction. 3D visualization of: (A) a binary mask representation of a nucleus segmented from a Fibroblast cell image; (B) a mesh representation of a reconstructed smooth surface of a nucleus; (C) three binary masks for nucleoli segmented within this nucleus; and (D) three mesh representations of nucleolar surfaces, color-coded along the *Z* axis. Visualizations are produced with the SOCR Dynamic Visualization Toolkit web application^57^.

### Morphometric measures

In this study, we used six shape measures as features quantifying geometric characteristics of the 3D surfaces, Fig. 1C. To calculate these measures, first the principal (min and max) curvatures (*к*_1_ ≤ *к*_2_) were computed using triangulated surface models representing the boundaries of genus zero solids^58^. Then, shape morphometry measures can be expressed in terms of principal curvatures: mean curvature as 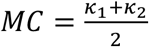, shape index as 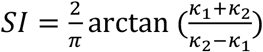, and curvedness as 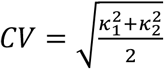. The principal curvatures of a surface are the eigenvalues of the Hessian matrix (second fundamental form), which solve for *k* |*H* − *kI*| = 0, where *I* is the identity matrix. If *S* is a surface with second fundamental form *H*(*X, Y*), *p* ∈ *M* is a fixed point, and we denote an orthonormal basis *u, v* of tangent vectors at *p*, then the principal curvatures are the eigenvalues of the symmetric Hessian matrix, 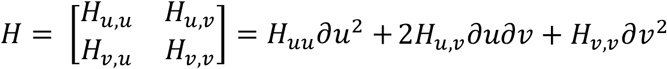, a.k.a. shape tensor. Let *r* = *r*(*u, v*) be a parameterization of the surface *S* ⊆ *R*^3^, representing a smooth vector valued function of two variables with partial derivatives with respect to *u* and *v* denoted by *r_u_* and *r_v_*, Fig. 3. Then, the Hessian coefficients *H_i,j_* at a given point (*p*) in the parametric *u, v*-plane are given by the projections of the second partial derivatives of *r* at that point onto the normal to *S*, 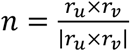, and can be computed using the dot product operator: *H_u,u_* = *r_u,u_* · *n, H_u,v_* = *H_v,u_* = *r_u,v_* · *n, H_v,v_* = *r_v,v_* · *n*, Fig. 3.

**Figure 3.**
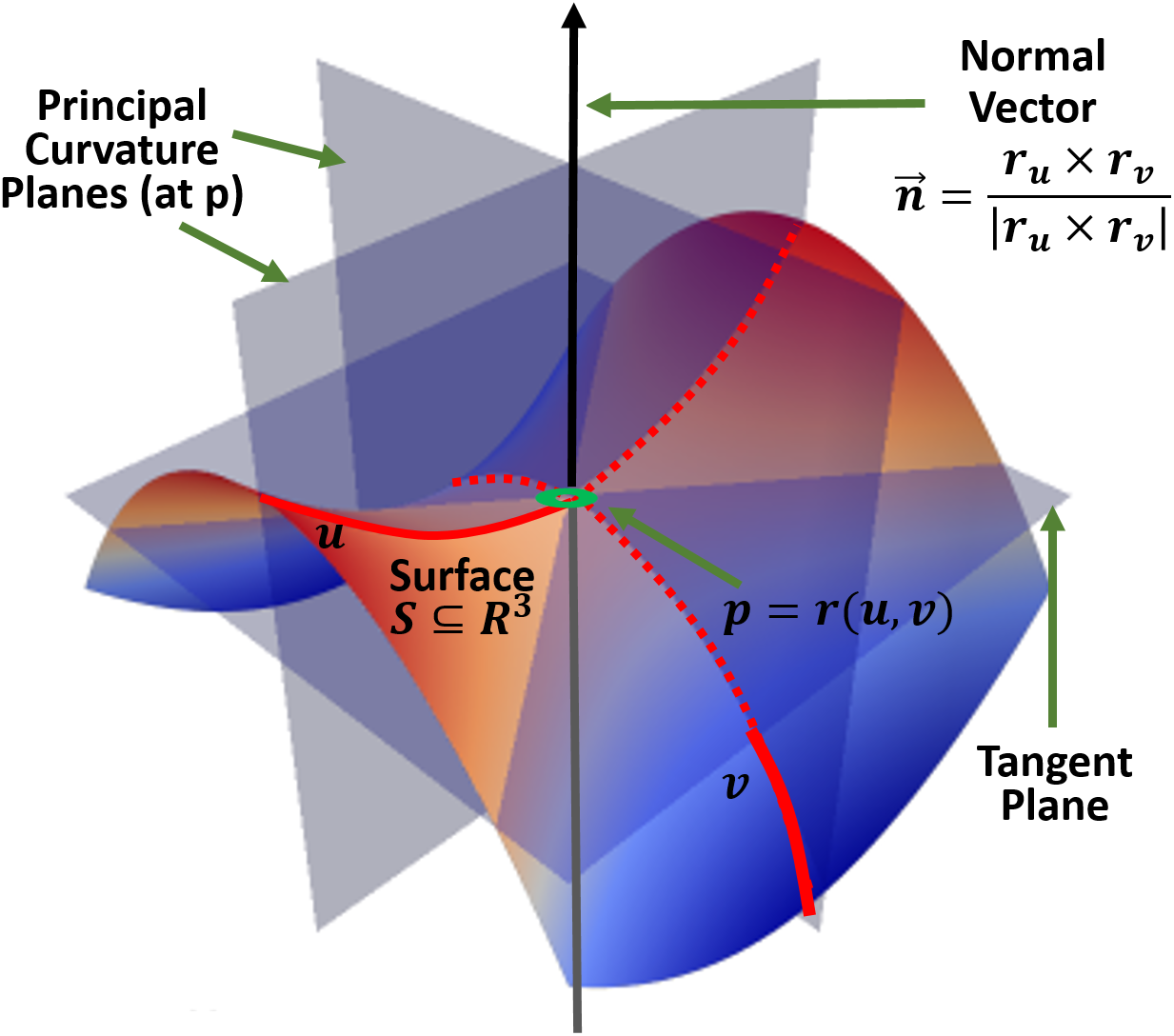
The (local) geometry of 2-manifolds. Per vertex definitions of curvature, relative to a local coordinate framework.

Volume is the amount of 3D space enclosed by a closed boundary surface and can be expressed as 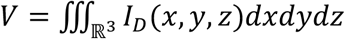, where *I_D_*(*x, y, z*) represents the indicator function of the region of interest (*D*)^59^. If *r*(*u, v*) is a continuously differentiable function and the normal vector to the surface over the appropriate region *D* in the parametric *u, v* plane is denoted by 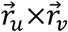, then *S_Ω_*:*r* = *r*(*u, v*), (*u, v*) ∈ *Ω*, is the parametric surface representation of the region boundary^60^. Then surface area can be expressed as 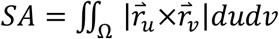. The fractal dimension calculations are based on the fractal scaling down ratio, ρ, and the number of replacement parts, *N*^61^. Accurate, discrete approximations of these metrics are used to compute them on mesh-represented surfaces^53,62^. These discrete metrics were first introduced as a part of the shape analysis protocol^31^ and were further applied in neuroimaging studies^32,33^.

The extracted 3D morphometric measures serve as features for training a number of machine learning algorithms in order to assess classification performance, Fig. 1D. The number of detected nucleoli per nucleus is included as an individual feature. We merged nucleoli-level features within each nucleus by computing sample statistics (e.g., average, minimum, maximum, and higher moments) for each morphometry measure^18^. These statistics are used to augment the signature feature vectors of the corresponding parent nuclei such that all feature vectors are of the same length. Correspondingly, nuclei that do not have any automatically detected internally positioned nucleoli were excluded from further analysis, such that for each nucleus there was at least one nucleolus.

### Visual analytics and machine learning for morphometric analysis

We performed exploratory visual analysis of extracted features using SOCRAT^49^, a web platform for interactive visual analytics. The goal of visual analytics is to support analytical reasoning and decision making with a combination of highly interactive visualizations and data analysis techniques^49,63^. SOCRAT implements a visual analytics workflow that encompasses an iterative process, in which data analysts can interactively interrogate extracted morphometric measures in the form of interactive dialogue supported by visualizations and data analysis components. In order to assess the variability of extracted morphometry data, we include t-Distributed Stochastic Neighbor Embedding (t-SNE)^64^ visualizations of the feature space generated by SOCRAT^49^. We also used SOCRAT to demonstrate interactions between the top-3 important features according to the best-performing classification algorithm. All derived morphometric datasets are made available within SOCRAT Web Demo application^65^. Finally, we demonstrated the ability to visualize volumetric images and extracted meshes online via SOCR Dynamic Visualization Toolkit web application^57^.

In general, correct classification of every single cell (type, stage, treatment, etc.) is a challenging task due to significant population heterogeneity of the observed cell phenotypes. For example, the same sample may contain a close mixture of intertwined “cancerous” and “non-cancerous” cell phenotypes; or, both classes may include apoptotic cells exhibiting similar shapes or sizes. Given the nature of cell samples, culturing, preparation, and collection, we have considered classification of cell sets rather than single cells. The idea of classifying sets of cells, rather than individual samples, is not new and has been used in recent biomedical image classification studies^12,66^. The rationale behind this is based upon the observation that even if an algorithm misclassifies a few cells in a sample, the final (cell set) label will still be assigned correctly, as long as majority of cells are classified correctly. Using this strategy, we performed classification on small groups of cells, ranging from 3 to 31 cells per set. During each fold of the internal cross-validation, these small cell sets are randomized by bootstrapping procedure with 1,000 repetitions. Random uniform sub-sampling is used to resolve the sample-size imbalance between the classes. Due to the possible presence of batch effects in data, we employed the Leave-2-Opposite-Groups-Out (L2OGO) cross-validation scheme^18^. L2OGO ensures that: (1) all masks derived from one image fall either in the training or testing set, and (2) the testing set always contains masks from 2 images of different classes. We used scikit-learn, a popular Python machine learning toolkit^67^, to evaluate a number of supervised classification algorithms.

### High-throughput workflow protocol

While the LONI Pipeline is a popular tool in neuroimaging and bioinformatics, it has been overlooked by the bioimage analysis community. In this work, we utilized the LONI Pipeline for the implementation of a streamlined multi-step protocol that relies on a diverse set of tools and solutions seamlessly connected in the LONI Pipeline workflow, Fig. 4. From a high-level perspective, every step of data processing and analysis protocol is wrapped as an individual module in the workflow that provides input and output specifications that allow the Pipeline to automatically connect and manage atomic modules. The modular structure of our implementation makes it highly flexible and not limited to specific tools included in the workflow. It can be repurposed for a wide range of different experiments by adjusting parameters, adding, removing, or replacing individual modules, while preserving high-throughput capabilities, as presented in the Discussion section. Every module represents an independent component that can be used in a stand-alone fashion. As a result, a distributed, massively parallel implementation of our protocol makes it possible to easily process thousands of nuclei and nucleoli simultaneously. The workflow does not depend on the total number of 3D objects, biological conditions, or a number of running instances since its execution is completely automated once the workflow configuration is fixed, including job scheduling and resource allocation. During the execution, our workflow provides a researcher with real-time information about progress and allows the viewing of intermediate results at every individual step. In addition, failed modules may easily be restarted.

**Figure 4.**
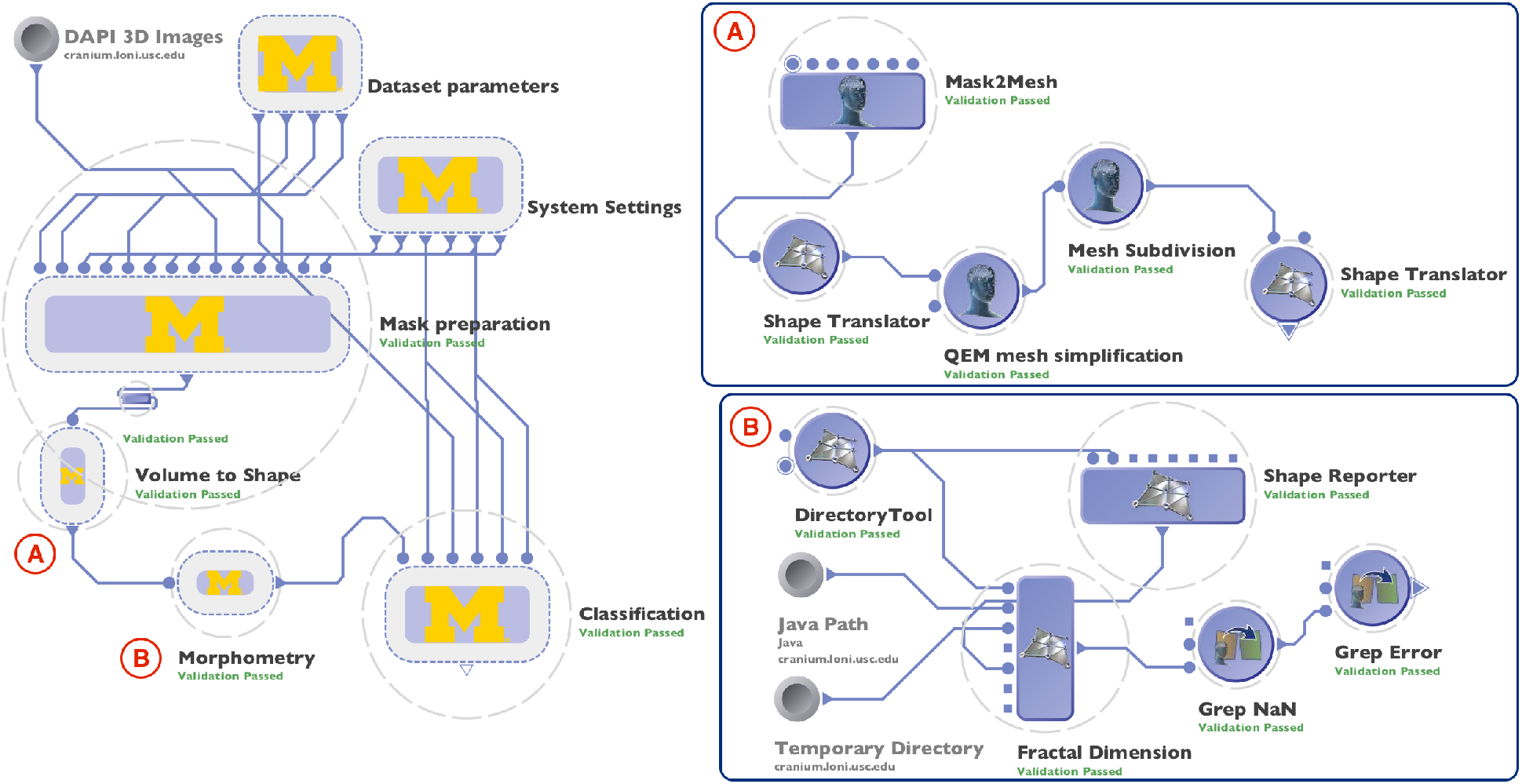
Screenshots of the exemplar graphical workflow in the LONI Pipeline client interface that include: (left) overview of the validated workflow protocol showing nested groups of modules; (A) expanded Volume to Shape group that includes modules that perform 3D shape modeling refinement; and (B) expanded Morphometry group that includes a module that performs morphological measure extraction.

The workflow is configured in such a way that it can consume data in the specific format we used, i.e. a 1024×1024×*Z* 3D volumes in different channels as 16-bit 3D TIFF files. Each volume is processed independently, in parallel fashion, such that workflow automatically defines how many processes are needed to analyze all of the input data. 3D shape modeling and morphometric feature extraction are performed on individual masks independently, which allows us to simultaneously run up to 1,200 jobs on the cluster during our experiments, effectively reducing the computing time. Finally, the workflow collects morphometry information from each individual mask and combines them in the results table that is further used as an input to classification algorithm. These capabilities allow the user to take advantage of modern computational resources, lift the burden of low-level configuration from researchers, make it easier to control the execution process, and improve reproducibility of the whole process.

## Results

### Validation on synthetic data

To validate the shape morphometry metrics, we first applied them to synthetically generated 3D masks. We used the scikit-image Python library^68^ to create 3D solids representing cubes, octahedra, spheres, ellipsoids, and 3 overlapping spheres with linearly aligned centers (Supplementary Fig. S1 online). We processed these objects and compare the resulting shape morphometry measures. Specifically, we aimed to confirm the expected close relation between the analytically derived measures of volume and surface area computed using the corresponding shape parameters (e.g., radius, size), and their computationally derived counterparts reported by the processing pipeline workflow. Our results illustrate that for nucleus-like shapes, e.g., spheres and ellipsoids, the computational error is within 2%. For faceted objects, e.g., cubes and octahedrons, the calculation error is within 6%. The increased error in the latter case can be explained by the mesh smoothing the surface reconstruction algorithm applies at the shape vertices to resolve points of singularity (e.g., smooth but non-differentiable surface boundaries).

To demonstrate the detection of shape differences between different types of 3D objects, we also compared overlapping spheres against circumscribed ellipsoids. As expected, the average mean curvature and curvedness measures are lower and shape index values are higher for spheres compared to ellipsoids. We observed a similar trend when comparing changes in these shape morphometry measures for spheres, ellipsoids, and overlapping spheres. For example, average mean curvature and curvedness were highest for overlapping spheres and lowest for spheres, which is expected based on definitions of these measures (see Supplementary Table S2). This simulation confirms our ability to accurately measure size and shape characteristics of 3D objects, which forms the basis for machine-learning based object classification based on boundary shapes. Exemplar results of synthetic data morphometry are available in Supplementary Table S1.

### Comparison with SPHARM for fibroblast nuclei shape classification

Classification of single cell nuclei from the fibroblast collection of the 3D Cell Nuclear Morphology Microscopy Imaging Dataset^18^ may be assessed using shape morphometry metrics as salient discriminatory features, which we did and compared against their corresponding SPHARM coefficients^16,25^. We used images of primary human fibroblast cells that were subjected to a G0/G1 Serum Starvation Protocol for cell cycle synchronization^69^. This protocol has previously been shown to alter nuclear organization, which was reflected in the observed morphology changes such as nuclear size and shape^70^. As a result, this collection contains 962 3D nuclear binary masks in the following phenotypic classes: (1) proliferating fibroblasts (PROLIF), and (2) cell cycle synchronized by the serum-starvation protocol cells (SS). We used these binary nuclear masks to calculate both SPHARM and morphometric features.

We obtained the SPHARM coefficients by using the popular SPHARM-MAT toolbox^71^ that implements surface reconstruction and spherical parametrization using the CALD algorithm^25^. Then, we followed by the expansion of the object surface into a complete set of spherical harmonic basis functions of degree *l* = 13 (default setting). Finally, SHREC method^72^ was used to minimize the mean square distance between corresponding surface parts. SPHARM shape descriptors were computed as described by Ducroz *et al*.^25^ and used as feature vectors for classification.

Throughout, we employed the open-source Python package scikit-learn 0.17.0^67^ to test a number of commonly used machine learning classification methods on derived feature vectors with default parameters for each method. Performance was compared using the L2OGO cross-validation scheme and the area under the receiver operating characteristic curve (AUC) as a performance metric. As shown in Table 1, 3D shape morphometric measures not only demonstrate comparable discriminative performance to SPHARM coefficients, but outperform them using all tested algorithms.

**Table 1.**
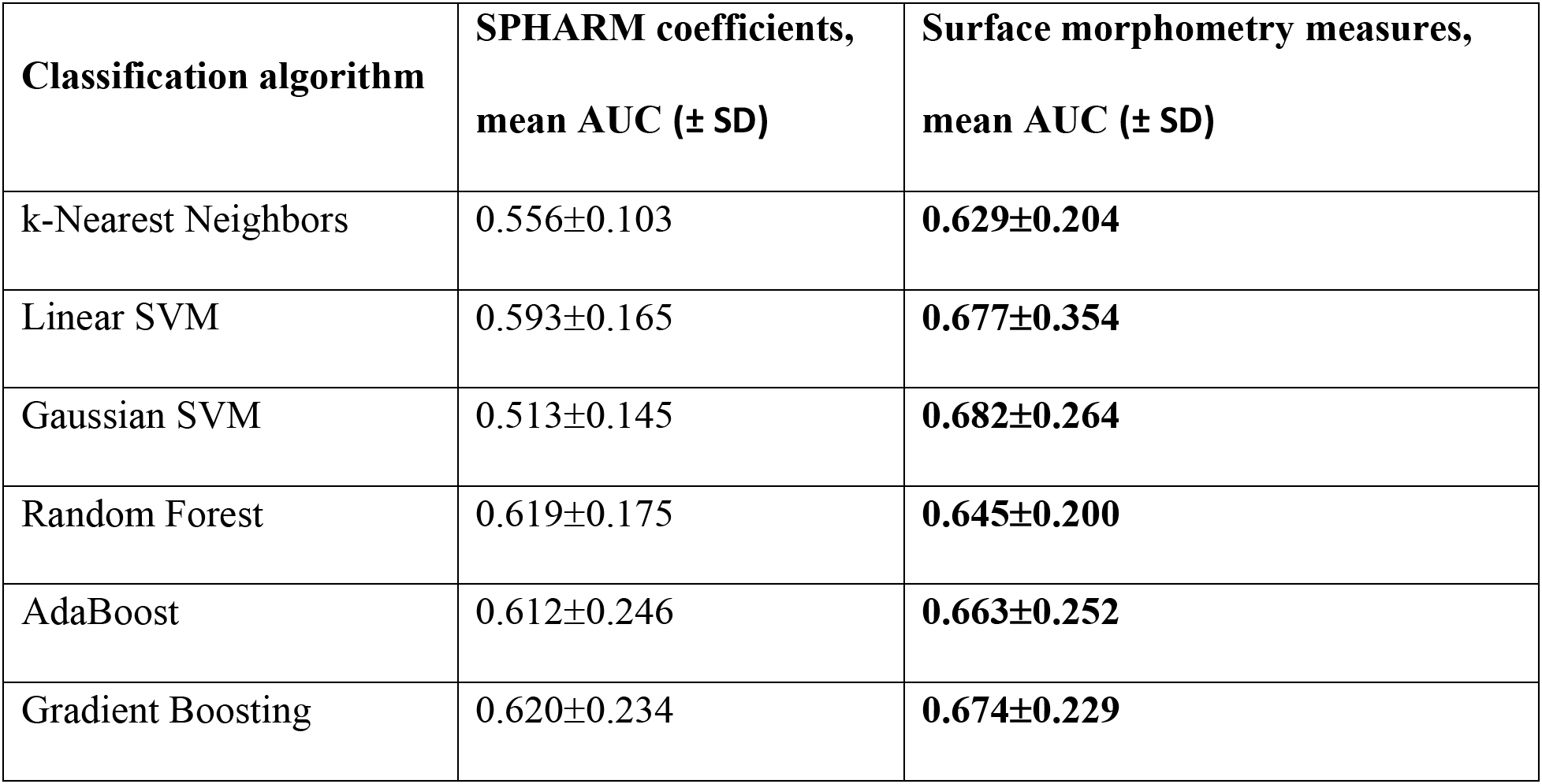
Comparison of SPHARM coefficients and our morphometry descriptors for single cell fibroblast nuclei classification

| **Classification algorithm** | **SPHARM coefficients,**<br>**mean AUC (± SD)** | **Surface morphometry measures,**<br>**mean AUC (± SD)** |
| --- | --- | --- |
| k-Nearest Neighbors | 0.556±0.103 | **0.629±0.204** |
| Linear SVM | 0.593±0.165 | **0.677±0.354** |
| Gaussian SVM | 0.513±0.145 | **0.682±0.264** |
| Random Forest | 0.619±0.175 | **0.645±0.200** |
| AdaBoost | 0.612±0.246 | **0.663±0.252** |
| Gradient Boosting | 0.620±0.234 | **0.674±0.229** |

### Fibroblast cell classification

The full collection of fibroblast masks for binary classification consists of total 965 nuclei (498 SS and 470 PROLIF) and 2,181 nucleoli (1,151 SS and 1,030 PROLIF). Figure 5A demonstrates the variability of the extracted morphometry measures in a t-SNE projection visualized in SOCRAT. Although there was a small degree of grouping, there was no clear separation between classes.

**Figure 5.**
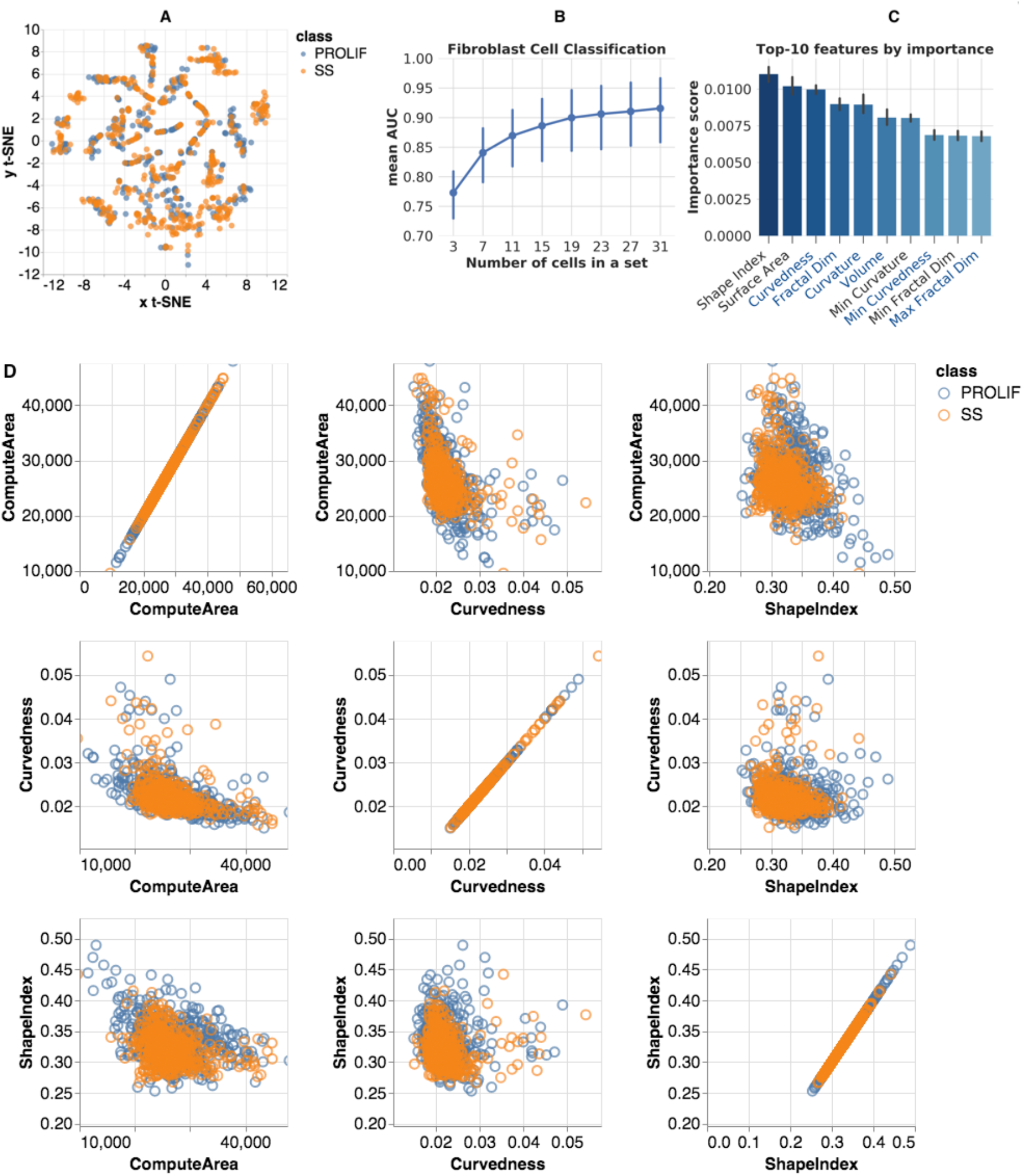
Fibroblast morphometric analysis: (A) SOCRAT visualization of t-SNE projection of morphometric feature space; (B) mean AUC for various cell set sizes; (C) top-10 features for classification by importance score (right, nucleolar feature names start with Avg, Min, Max or Var, feature names that were also reported in top-10 for PC3 cells are shown in blue font); and (D): SOCRAT visualization of interactions between top-3 features.

The best result by a single classifier was achieved using a stochastic gradient boosting classifier with 1,500 base learners, maximum tree depth 8, subsampling rate 0.5. Hyper-parameters were fine-tuned using a cross-validated grid search. To evaluate these classification results, we measured accuracy, precision, sensitivity, and AUC over L2OGO cross-validation, which are presented in Table 2 for single cell and 19-cell-set classifications. Figure 5B shows mean AUC values for set sizes from 3 to 19 cells. A 90% mean AUC was reached when classifying sets with 19 cells and 92.5% for sets with 31 or more cells.

**Table 2.** Fibroblast single cell and 9-cell sets classification accuracy

| **Measure** | **Single cell,**<br>**mean (± SD)** | **19 cells set, mean (± SD)** |
| --- | --- | --- |
| **Accuracy** | 0.699 (±0.076) | 0.899 (± 0.123) |
| **Precision** | 0.701 (±0.075) | 0.922 (± 0.115) |
| **Sensitivity** | 0.692 (±0.127) | 0.874 (± 0.224) |
| **AUC** | 0.699 (± 0.076) | 0.899 (± 0.123) |

The gradient boosting classifier also computes and reports cross-validated feature importance (Fig. 5C). These allow us to evaluate which measures differ between two cell conditions, and potentially propose novel research hypotheses that can be tested using prospective data. Previous analysis has reported quantifiable changes in both nuclear size and shape under serum-starvation^70^. In our results, both nuclear (top-6, out of top-10) and nucleolar (4 of top-10) morphometric size and shape features are reported to be of high importance for distinguishing SS fibroblasts from PROLIF (Fig. 5C). We also visualized the relationship between top-3 features using SOCRAT, see Fig. 5D. Visualizations suggest the smaller variation of morphometric measures in SS fibroblast nuclei compared to their PROLIF counterparts. This result may provide insight in further downstream analysis of potential underlying mechanisms that lead to these morphometric changes. We made the fibroblast morphometry data publicly available within SOCRAT for further analysis and validation^65^.

### PC3 EPI/EMT cell classification

The second collection contains images of human prostate cancer cells (PC3). Through the course of progression to metastasis, malignant cancer cells undergo a series of reversible transitions between intermediate phenotypic states bounded by pure epithelium and pure mesenchyme^9^. These transitions in prostate cancer are associated with quantifiable changes in both nuclear and nucleolar structure^10,73^. PC3 cells were cultured in: (1) epithelial (EPI), and (2) mesenchymal transition (EMT) phenotypic states. The collection includes 458 nuclear (310 EPI and 148 EMT) and 1,101 nucleolar (649 EPI and 452 EMT) 3D binary masks. Figure 6A demonstrates the variability of the extracted morphometry measures in a t-SNE projection visualized in SOCRAT. Similar to fibroblasts, the projection of the PC3 morphometric feature space does not demonstrate clear separation between classes.

**Figure 6.**
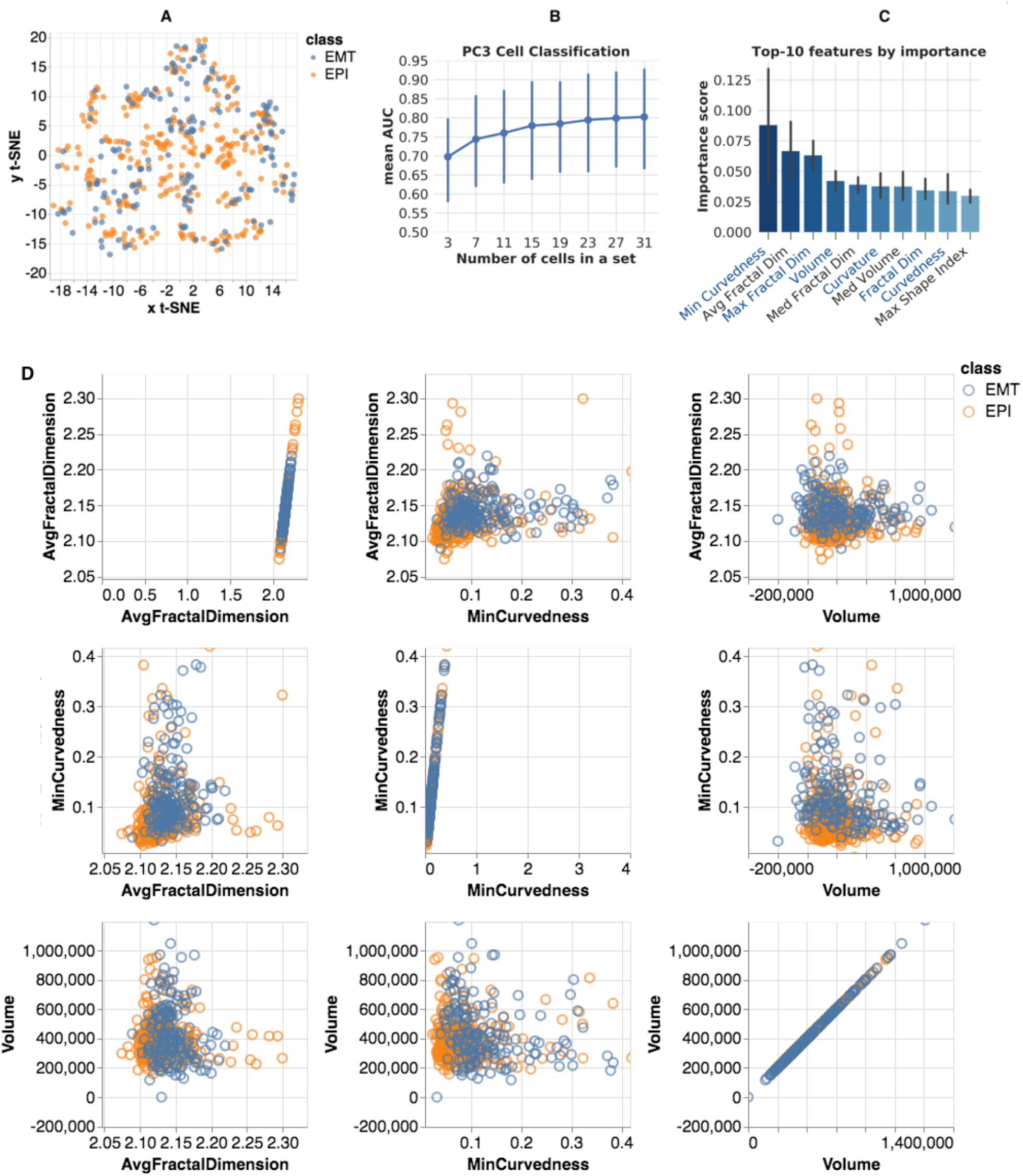
PC3 morphometric analysis: (A) SOCRAT visualization of t-SNE projection of morphometric feature space; (B) mean AUC for various cell set sizes; (C) top-10 features for classification by importance score (right, nucleolar feature names start with Avg, Min, Max or Var, feature names that were also reported in top-10 for Fibroblast cells are shown in blue font); and (D): SOCRAT visualization of interactions between top-3 features.

In this case, the best classification performance by single classifier is the result of applying a random forest model (1,000 trees, maximum tree depth 12, maximum number of features for the best split 40%). Hyper-parameters fine-tuning, accuracy metrics, and cross-validation procedures are identical to the ones reported in the previous fibroblast experiment. Classification of sets of 19 cells achieves a mean AUC of 76.2%, Table 3. Figure 6B reports the AUC for different group sizes to show how the classification performance increases with the cell-set size and reaches 80% for sets of 27 or more cells. In this experiment, we also examined the classifier-reported feature importance, Fig. 6C. The top-10 important features in this classification included nuclear (4 of top-10, which were also in Fibroblast top-6) and nucleolar (top-3, 6 out of top-10) shape morphometry features. Top feature interactions visualized using SOCRAT demonstrate the important changes in distributions of nucleolar morphometric measures, Fig. 6D. For example, it seems that the EPI nucleoli tend to have more variability in minimal curvedness and average fractal dimension, compared to EMT nucleoli. Previously reported PC3 morphological analyses^73^ only used simple 2D nuclear form measures, such as diameter and the size of the bounding box. While we confirmed the importance of nuclear form in our results and suggested the need for further investigation of other highly ranked features, such as nucleolar curvedness, shape index, and fractal dimension, which may provide additional mechanistic insights. PC3 morphometry data are made publicly available within SOCRAT for further analysis and validation^49^.

**Table 3.**
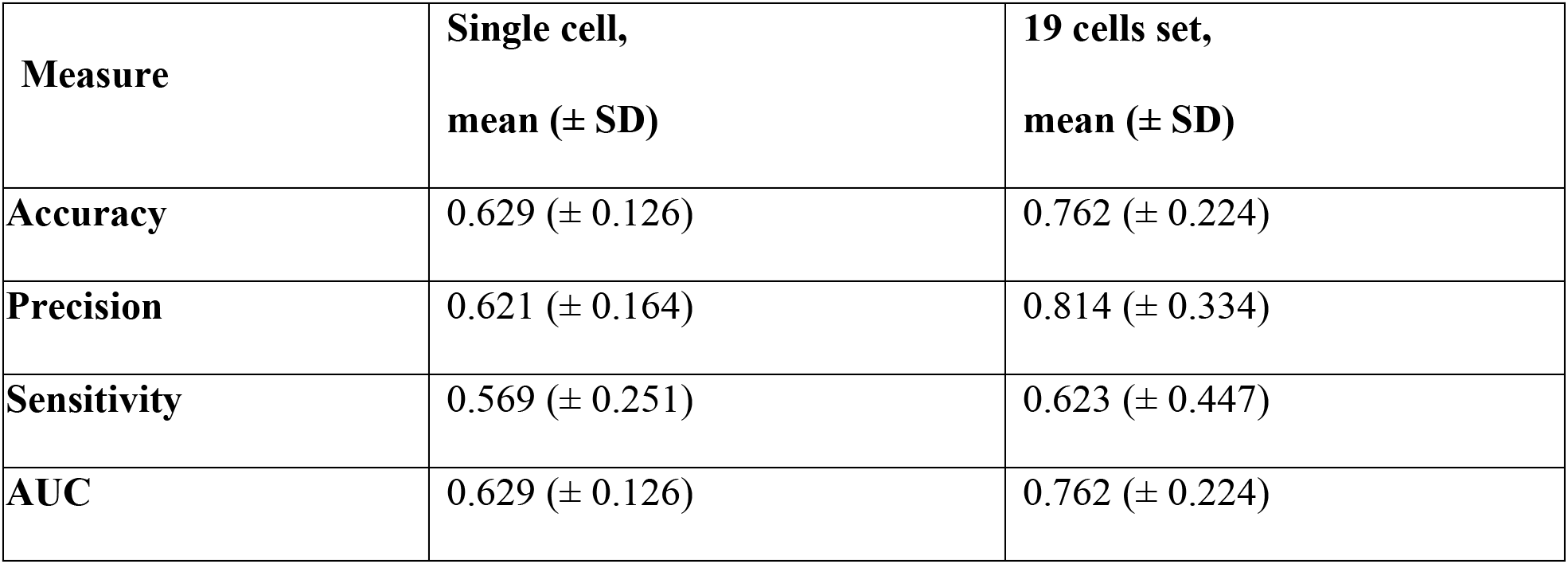
PC3 single cell and 9-cell sets classification accuracy

## Discussion

In this study, we proposed, implemented, and validated a solution for 3D modeling, morphological feature extraction, analysis, and classification of cells by treatment conditions. Compared to other studies using 2D projections, this approach operates natively in 3D space and takes advantage of extrinsic and intrinsic morphometric measures that are more representative of the real, underlying nuclear and nucleolar geometry and allow easy human interpretation. Given the limitations of using 3D voxels for accurate shape representation, we employed 3D surface models to extract more informative size and shape measures to improve the morphology classification performance. Robust surface reconstruction allows accurate approximation of 3D object boundaries that was validated on synthetic data. Suggested shape morphometric measures outperform another popular approach and demonstrated their universality across different cell types, conditions, and even domains^31–33^.

Our computational protocol implementation is highly parallel with throughput, limited only by the number of available computing nodes, and it can process thousands of objects simultaneously with minimal human intervention. This pipeline workflow integrates a number of open-source tools for different steps of data processing and analytics. Every module in our workflow represents an individual component that can be easily modified, removed, or replaced by an alternative. Such modular software platform architectures have been shown to enable high reusability and ease of modification^49^. This allows the user to use the same workflow or customize and expand it (e.g., specification of new datasets, swapping of specific atomic modules) for other purposes that require the analysis of a diverse array of cellular, nuclear, or other studies. The live demo available via the LONI Pipeline demonstrates the simplicity of use and high efficiency of parallel data processing. LONI also provides guest access (see Supplementary Information) and an opportunity to utilize a 4,500-core LONI cluster after applying for a collaboration account.

We tested our approach on the 3D Cell Nuclear Morphology Microscopy Imaging Dataset^18^, which includes a total of ~1,500 nuclear and ~2700 nucleolar masks. The classification results on these data comparing epithelial vs. mesenchymal human prostate cancer cell lines, and serum-starved vs. proliferating fibroblast cell lines, demonstrate the high accuracy of cell type prediction using 3D morphometry, especially when applied to sets of cells. Although different classification algorithms appear to be optimal for different experiments, we observed that both nuclear and nucleolar morphometric measures are important features for discriminating between treatment conditions or cell phenotypes. In the case of fibroblast classification, the results show the importance of nuclear morphometry, the number of nucleoli per nucleus, and various internal nucleolar morphometric measures. These observations confirm and extend previously reported results. For PC3 cells, the most important classification features are the moments of the distributions of various nucleolar morphometric measures, along with nuclear size and shape. Interestingly, there were 3 common morphometric features among the top-10 most important ones for both cell lines. This confirms previously reported observations^73^, suggests new important morphological characteristics, and demonstrates that our method extracts relevant information from cell forms to successfully classify cells using a combination of criteria. In addition, this also demonstrates the importance of sophisticated shape metrics, compared to volume and surface area, that alone, were not the most informative features for the classification results. The use of SOCRAT enables interactive interrogation of morphometric data in a visual manner, supported by analytical tools. This method of interactive ‘visual analytics’ provides insight into feature dependencies and interactions, and can be used for result interpretation. We also demonstrated the visualization of 3D volumetric images and derived meshed surface representations using the SOCR Dynamic Visualization Toolkit web application^57^.

Our computational approach is scalable and capable of processing complex big 3D imaging data, and is not limited to nuclear and nucleolar shapes. With some changes, it can be applied to other cellular and nuclear compartments of interest. More specifically, the robust smooth surface reconstruction algorithm can be directly applied to any 3D shapes, as long as their topology is sphere-like. Together with molecular level techniques, such as Hi-C, our 3D shape morphometry workflow can form a powerful combination for the investigation of DNA architecture in the spatial and temporal framework of the 4D nucleome^6,74^. One example of the many possible future applications of this workflow is to study asymmetric cell division. Stem and progenitor cells are characterized by their ability to self-renew and produce differentiated progeny. A balance between these processes is achieved through controlled asymmetric divisions and is necessary to generate cellular diversity during development and to maintain adult tissue homeostasis. Disruption of this balance may result in premature depletion of the stem/progenitor cell pool, or abnormal growth^75,76^. In many tissues, dysregulated asymmetric divisions are associated with cancer. Whether there is a causal relationship between asymmetric cell division defects and cancer initiation is unknown. We proposed that our shape analysis pipeline will be useful in studying the 4D nucleome topology of morphogenesis and cancer initiation.

As one of the approach limitations, we pointed out that other geometric measures can be used to characterize shapes of interest, such as intrinsic shape context, compactness, symmetry, smoothness, convexity, etc. In the current representation, analyzable shapes are limited to genus zero surfaces, which is a fair assumption when modeling objects like nuclei or nucleoli. However, it might be not trivial when considering other nuclear structures, for example, chromosome territories or interchromosomal loops, since their topologies may not be homeomorphic to a sphere, or may not appear to be genus zero under some imaging conditions and modalities. It is also conceivable, yet not very likely for the discretized LB, that 2 different shapes may have the same spectra. In this case, the algorithm may fail to detect the intrinsic differences between them due to false-negative error. Even though our workflow only requires little intervention (classifier selection and tuning), further improvements would involve adaptive implementations with even less manual intervention, as well as extraction of additional features. Another option is to use deep learning-based methods that alleviate the need to define features and allow to learn relevant patterns directly from data^77,78^. Recent applications have demonstrated the ability of deep neural networks to successfully perform classification, segmentation, and detection on limited amounts of biomedical imaging data^79–81^. For example, textural features could possibly increase discriminatory power of the method and provide more information on chromatin reorganization^82^. Since nuclear deformation serves as a proxy to underlying processes, the importance of particular features and the method’s ability to classify nuclei does not provide direct insight into the fundamental biological mechanism driving the observed morphometric differences between cell phenotypes or environmental conditions. The computational results should be further tested and externally validated using other experimental conditions and prospective data.

## Conclusions

Quantification of cell nuclear morphology enables more subtle characterization of cellular phenotypic traits, which can be associated with functional changes coupled to underlying biological processes. Using the new methodology described in this paper, we compared the morphology of serum-starved vs. proliferating fibroblast cells as a control, followed by a comparison of epithelial with mesenchymal human prostate cancer cell lines. In the case of fibroblast classification, our results show the importance of nuclear morphometric change, along with the number of detected nucleoli per nucleus, and various internal nucleolar morphometric measures. Results for PC3 cells demonstrate that the changes in nucleolar morphology are the most informative. However, in both cell lines, both nuclear and nucleolar morphometric measures contribute to the discriminative power of the classification algorithms. To the best of our knowledge, this study is the first where a 3D morphometric assay could easily distinguish between the epithelial and mesenchymal cell nuclei. These findings suggest that further investigation of highly ranked features that were not previously reported, such as nucleolar curvedness, shape index, and fractal dimension, may provide interesting mechanistic insights.

The ability to automate the processes of specimen collection, image acquisition, data preprocessing, computation of derived biomarkers, modeling, classification, and analysis can significantly impact clinical decision-making and fundamental investigation of cell deformation. To our knowledge, this is the first attempt to combine 3D cell nuclear shape modeling by robust smooth surface reconstruction and extraction of shape morphometry measures into a highly parallel pipeline workflow protocol for morphological analysis of thousands of nuclei and nucleoli in 3D. This approach allows efficient and informative evaluation of cell shapes in the imaging data and represents a reproducible technique that can be validated, modified, and repurposed by the biomedical community. This facilitates result reproducibility, collaborative method validation, and broad knowledge dissemination.

## Availability of materials and data

The documentation supporting the conclusions of this article together with the pipeline workflows and underlying source code are made available online on the project webpage SOCR 3D Cell Morphometry Project, http://socr.umich.edu/projects/3d-cell-morphometry^54^.

## Author contributions

AA and DD designed and built modules for data pre-processing and curation. AAK, AA, DD, AAF, GVF, WM, JRD, GAH, GZ, AC, JWW iteratively evaluated various image analysis approaches. AAK, AAF, SSH and IDD designed and built modules for 3D shape modeling, morphometric feature extraction, and visual analytics. AAK, AAF, AA, and IDD designed, built, and executed high-throughput Pipeline workflows. AAK implemented and performed statistical analysis and result interpretation. GVF, WM, JRD, GAH, GZ, AC, JWW, JEV, RWV, KJP and DSC contributed to conceptualization of study. IDD and BDA conceived the study. AAK, AAF, AA, IDD and BDA wrote the manuscript with input from other authors. All authors participated in numerous project discussions and decision making. All authors read and approved the final manuscript.

## Acknowledgements

Many colleagues at the Michigan Institute for Data Science (MIDAS), the Department of Computational Medicine and Bioinformatics, and the Statistics Online Computational Resource (SOCR) provided contributions including ideas, pilot testing, improvement suggestions and other assistance in the development and validation of these methods. We also would like to thank colleagues at the Laboratory of Neuro Imaging (LONI), at the Keck School of Medicine, University of Southern California, for providing technical support for the LONI Pipeline environment.

